# CSTDB: A Crop Stress-tolerance Gene and Protein Database Integrated by Convolutional Neural Networks

**DOI:** 10.1101/456343

**Authors:** Di Zhang, Yi Yue, Yang Zhao, Chao Wang, Xi Cheng, Ying Wu, Guohua Fan, Panrong Wu, Yujia Gao, Youhua Zhang, Yunzhi Wu

## Abstract

Numerous studies have shown that many genes and proteins in plants are involved in the regulation of plant resistance to abiotic and biotic stresses. The researches on the stress tolerance of crops are also the focus of many researchers. To provides a reliable platform for collecting and retrieving genetic and protein information related to stress tolerance found in crops, we constructed CSTDB(Crops Stress-tolerance Database), an integrated database that includes stress-tolerance genes and proteins for many crop species. The database was developed based on convolutional neural network technology. It is a web-accessible database that contains detailed information on the stress-tolerance genes and proteins of major crop species. Currently, the database records four major crops containing 1,371 abiotic stress-tolerance genes or proteins, and 207 genes or proteins associated with biotic stress. Each gene and protein has detailed functional information and sequence information, such as stress types, Genbank ID, Pubmed ID, Protein ID, 3D model picture and FASTA files. As a user-friendly browsing tool, this database provides search functions, BALST functions and file download functions. CSTDB can be a valuable resource, which is designed to meet the broad needs of researchers working on crops stress-tolerance experiments. Database URL: http://pcsb.ahau.edu.cn:8080/CSTDB

## Background

In nowadays society, crops are still the guarantee for the survival of all humanity. They provide a significant proportion of the calories consumed by humanity so maintaining and improving upon current production levels will be critical to provide food security for a growing world population(Pearce *et al.* 2015). With the continuous development of agricultural modernization, the output of crops is also increasing. However, due to the increasing population of the world, the storage of crops is still in short supply(Dubois *et al.* 2011). According to current trends, the gap between population and food supply is widening(Muazu *et al.* 2017). The reason for this phenomenon is that various crops are affected by the external environment (drought(Magwanga *et al.* 2018), heat(Hassen *et al.* 2018), frost(Kruse *et al.* 2017), salinity(Wang *et al.* 2016), waterlogging(Valliyodan *et al.* 2016)) and plant pathogens, and they suffer varying degrees of damage from seed germination to growth and maturation. According to the form of destruction, we classify these damages as abiotic (Erpen *et al.* 2017) and biotic stresses (Borrill *et al.* 2017). Among them, drought, salinity and waterlogging are the big three abiotic stresses that limit crop yields, resulting in a large number of losses worldwide(Zhang *et al.* 2017).In this background, numerous researchers are engaged in the study of crop stress-tolerance and have many achievements, such as breeding stress-tolerance varieties by hybridization techniques(Rashid *et al.* 2017; Marcińska *et al.* 2018), obtaining genetically-resistant varieties(Shivakumara *et al.* 2017; Wang *et al.* 2016). There are also many researchers who have discovered biological genes that play a positive role in resisting stress in plants(Shi *et al.* 2017; Li *et al.* 2015).

Rice(Oryza sativa), barley(Hordeum vulgare), wheat(Triticum aestivum), and corn(Zea mays) are all important crops in the world, and there are plenty of planting areas in the world (Chung *et al.* 2011; Sacks *et al.* 2016). However, due to abiotic stress and biotic stress, the yield of these crops is often worrying(Bray *et al.* 2000). Over the past few decades, with the continuous development of biological technologies and high-throughput sequencing technologies(Goff *et al.* 2002; Yu *et al.* 2002; International Barley Genome Sequencing Consortium *et al.* 2012; International Wheat Genome Sequencing Consortium *et al.* 2014; Schnable *et al.* 2009), more and more researchers have been able to identify and clone genes or transcription factors related to stress resistance in these crops(Mickelbart *et al.* 2015; Wang *et al.* 2016). As a large number of genes and transcription factors were found to be related to the stress resistance of crops(Güre *et al.* 2016;Barnawal *et al.* 2017; Wang *et al.* 2017; Gao *et al.* 2018),researchers published the research results through the display of articles in major academic journals for reference by other researchers. Mass research were scattered across different journals’ websites or public document data websites. Hence, it is very significant and meaningful to collect and integrate the information in these articles into a single database to present various information related to stress-tolerance in different species, and to further clarify the mechanisms of stress and regulation. Under these conditions, several database websites on plant (such as Arabidopsis, rice, etc.) stress-tolerance genes have been released in public, such as STIFDB(Shameer *et al.* 2009), STIFDB2(Naika *et al.* 2013), DroughtDB(Alter *et al.* 2015), RiceSRTFDB(Priya *et al.* 2013)and PASmiR(Zhang *et al.* 2013). These database analyzed and integrated the collected genetic sequence data or literature data through different methods, and finally established website for researchers who are engaged in related research. The screening of these data is the result of manual screening, which requires a lot of manpower and time. At the same time, some researchers also use computer-based machine learning techniques to process and integrate the collected data. For example, the PPIM database is constructed using the SVM technology to predict the corn protein interaction network(Zhu *et al.* 2015), and Payam Karisani et al. proposed a biomedical data set retrieval method based on probability and machine learning (Karisani *et al.* 2018).

Based on this background, our experiment used deep learning techniques to classify a large number of literatures' abstracts related to rice, barley, wheat, and corn collected from public data sites. We used unsupervised methods to let computer automatically identify whether the literature is related to crops stress-tolerance genes. According to the results after the screening, we further extracted the detailed information of the corresponding literature, including the species name, type of stress, gene name, PubMed ID of literature, 3D model(Waterhouse *et al.* 2018) and so on. Finally, based on all the data we have integrated, we set up a database for stress-tolerance genes of some crops. We provide an excellent information platform for researchers working on related work.

## MATERIALS AND METHODS

### Data sources

All data for this data is obtained from NBCI website. According to the keyword resistance of major crops provided by National Joint Engineering Laboratory of crop resistance breeding and disaster reduction(Supplementary Table S1), we used these keywords as search conditions, and used the crawler technology to collect the abstracts of related literatures on the NCBI website based on the NCBI Web Services interface. In the end, we collected 33,207 abstracts on the NBCI website. According to the collected data, the literature on rice has the most abstracts, reaching 16,105, of which the most research on rice blast has reached 7,870. The most research of barley and wheat is the effect of stripe rust on its growth, and the most stress on corn is heat. These collected abstracts of literatures are the data sets for this study. An overview of these data is shown in Fig. 1.

**Figure 1.**
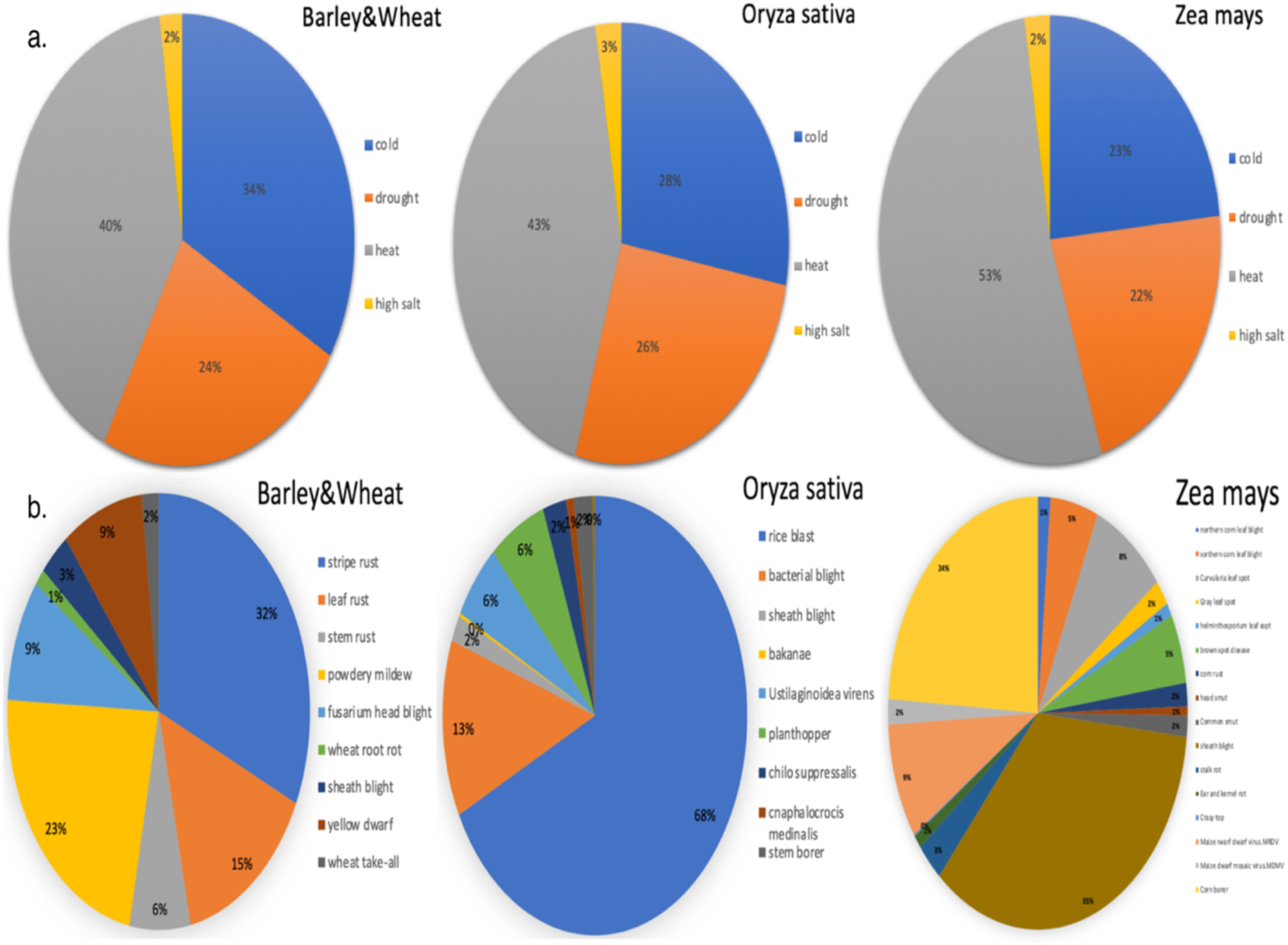
a Proportion of dataset types of abiotic stress tolerance for different crops. b Proportion of dataset types of biotic stress tolerance for different crops

### Data classification and integration

We first preprocessed the acquired documents, including data cleaning and word segmentation, and then construct the word vector based on Word2vec and extract its features (Goldberg *et al.* 2014). The processed word vector were then embedded into the Eembedding layer of TextCNN(Kim *et al.* 2014), and the TextCNN model were used to train it using the training set. Finally, we used the test set to test the trained model. After the test results meet the requirements, we finally classified and predicted the remaining data. The specific parameters of the experimental equipment are shown in the Table. 1. The workflow of data classification and integration is shown in Fig. 2.

**Table 1.** Parameters of the experimental equipment

| Experimental configuration | Environment configuration |
| --- | --- |
| Operating system | Ubuntu16.04 |
| CPU | Inter core i7 -4790 3.60GHZ |
| Programming language | python3.0 |
| Deep learning framework | TensorFlow1.4.0 |

**Figure 2.**
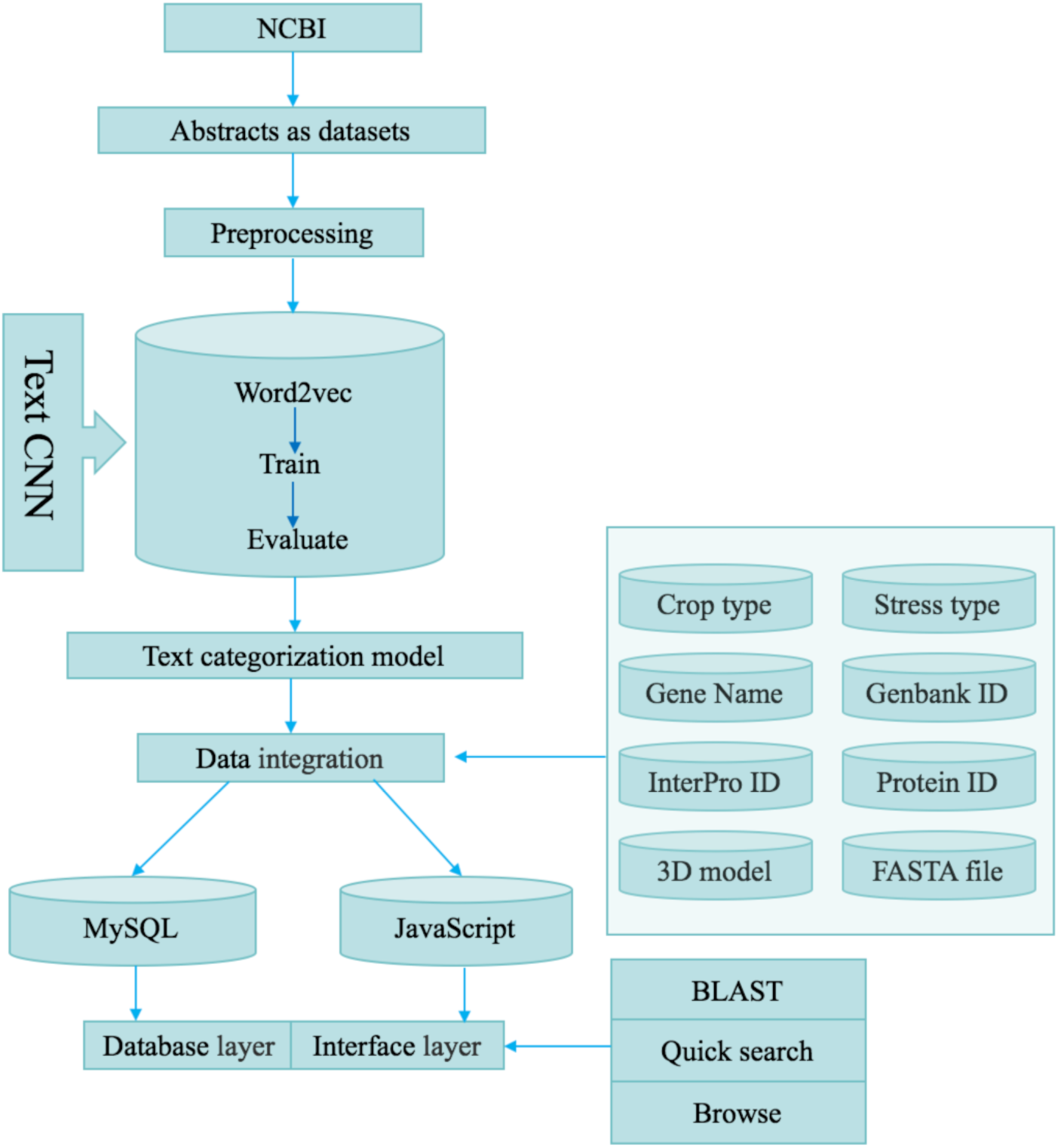
The workflow of data classification and integration

Through this classification model, a total of 1821 abstracts related to crop stress-tolerance genes and proteins were identified. Subsequent to the false positive removal of these abstracts, a total of 1088 target literature abstracts were screened out. After browsing the full text of the literature, 1240 target genes and proteins were extracted. In terms of the species of crops, rice had the highest number of a stress-tolerance genes and proteins, 803, followed by barley and wheat with 567 and corn with 208. According to the category of stress, there are 1,371 genes related to biotic stress, and 207 genes related to abiotic stress. The number of stress-tolerance genes and proteins of different crops is shown in Figure 3.

**Figure 3.**
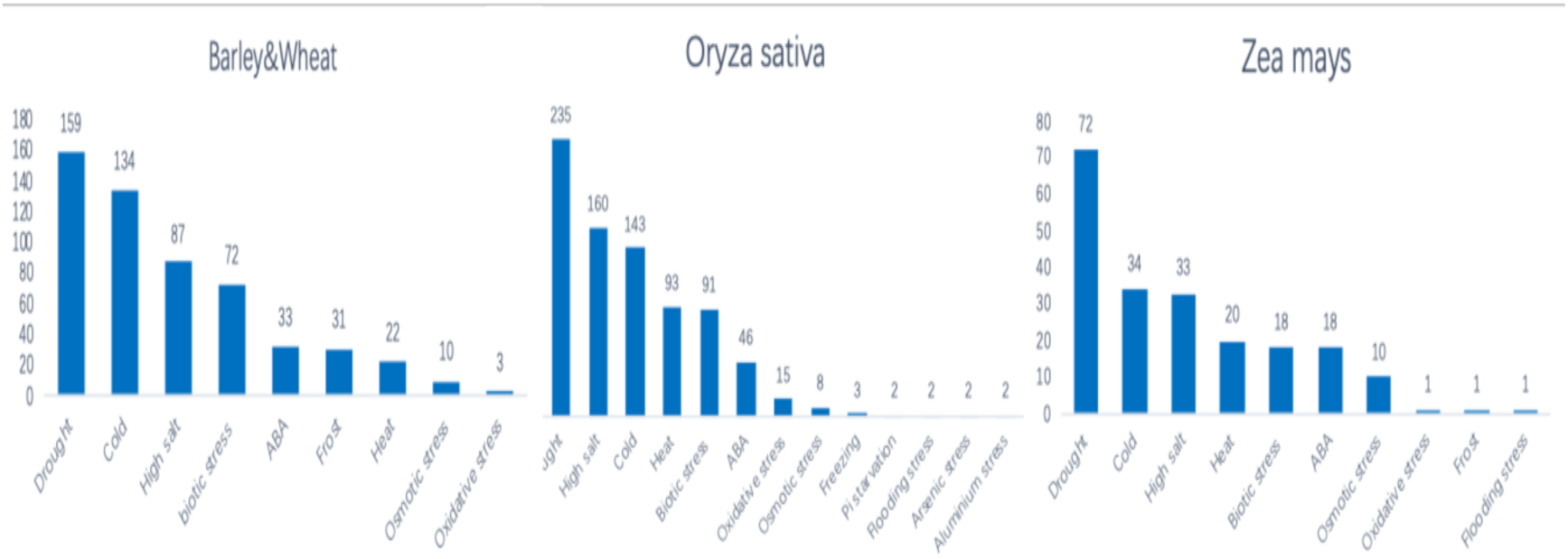
The number of stress-tolerance genes and proteins of different crops

### Website implementation

The webite was constructed using several different programming packages. The user interface of CSTDB was developed on framework of HTML (Cohen *et al.* 2002) and JSP (Kurniawan *et al.* 2002). It can make the website automatically adjust and adapt to different device screen sizes. The back-end of website was implement by springMVC(Zhang *et al.* 2013), Hibernate(Bauer *et al.* 2005) and MySQL(Welling *et al.* 2003).

### Database implementation

The database implementation uses a flexible storage schema to store the data. It was maintained in the Windows server 2003 equipped with a 2 GHz CPU processor and 2 GB of RAM. There is a Java application in back-end that uses Apache Tomcat server 7.0 technologies such as Java server Pages (JSP) and Java servlets to manage web-based data input and data queries.

## RESULTS AND DISCUSSION

### Web interface

This website provides an intuitive and friendly graphical interface that can be adapted to different devices through the layout of the web page (Fig. 4b), thus providing users with the most comfortable browsing experience. The results displayed on all web pages are presented as separate windows containing detailed information on the target gene or protein, such as the species, gene name, Genbank ID, PubMed ID, 3D model, FASTA file and so on(Fig. 4e). On the home page, the website provides users with three browsing modes: all, by species type, and by stress type(Fig. 4a,4d). Users can view all the stress-tolerance genes and proteins of a single crop, or you can obtain all the genes and proteins related to one stress by selecting the name of the stress in the sub-list for accurate screening.

**Figure 4.**
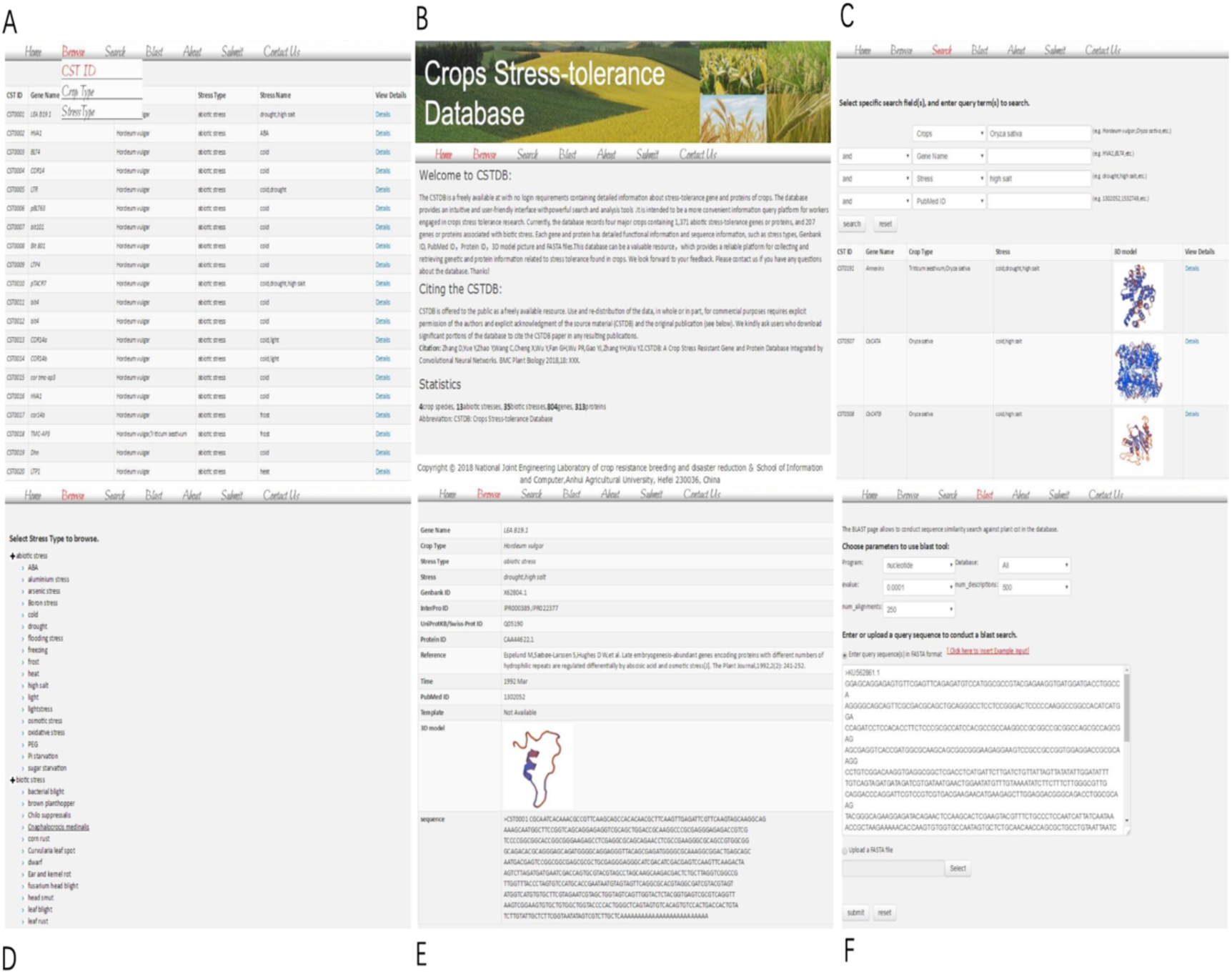
Web interface of the CSTDB

### Search module

The CSTDB provides a full-featured search function(Fig. 4c). The users can perform fuzzy search or precise search on the database according to his own requirements. They can input the name of the crop in the search field to obtain the stress-tolerance genes and proteins of all the crops. At the same time, users can also enter the gene name, Genbank ID, PubMed ID or stress name for precise search.

## BLAST

In order to achieve the sequence similarity search function, the database also provides users with a homology search tool 'BLAST'(Fig. 4f). This tool used a stable version of NCBI BLAST-2.7.1 + as a back-end BLAST executable to embed in the back end of the database, while using the SequenceServer app as the front-end input(Zhu *et al.* 2017; Camacho *et al.* 2009). This website provides a more intuitive graphical interface for this function, making it easy and fast for users to use BLAST search.

### Submit

The CSTDB provides a submit page that allows other researchers to independently contribute new reliable information for crops stress-tolerance genes or proteins as they become available. For user submissions, primary required fields are: 1) Crop species, 2) Stress type, 3) Stress name, 4) Gene name, 5) Genbank ID, 6) Protein ID, 7) 3D model, 8)Reference , 9) PubMed ID. The curator at our site conducts manual verification of the original publication for data validation purposes to maintain the quality and integrity of the database. Submissions that pass this review process are then approved for entry into the CSTDB database. The all new submission will be made available in coming CSTDB versions on a monthly release schedule.

### Future work

Our database is based on NCBI's existing literature abstracts prior to May 2018 as datasets. As the research literature on the NCBI website continues to increase, we will update the database on a regular basis. The current database collects information on four crops, and in the next study we will increase the types of crops, such as Soybean and Peanut. We will also provide a user submission function for the website. Users can upload relevant documents found in the research to our database by filling in the blanks, which will help the database become more comprehensive.

### Conclusions

We built CSTDB based on the NCBI website using convolutional neural network technology, which is a database integrating stress-tolerance genes and proteins of several crops. The database provides an intuitive and user-friendly interface with free access to browsing information and downloading data. The database integrates multiple information on a variety of crop stress-tolerance genes and proteins, and provides powerful search and analysis tools that allow users to quickly identify and analyze target genes or proteins. This database provides a more convenient information query platform for workers engaged in crops stress tolerance research. In addition, our data processing method provides an effective reference value for the construction of related databases in the future. In the next study, we will continue to supplement this database by adding more species and more detailed information on the genes and proteins to this database, so that it can provide researchers with more quality services.

## Acknowledgments

This research was supported by National key research and development project of China (No. 2017YFD0301303),National Natural Science Foundation of China (No.31671589),the Anhui Agricultural University High-level Scientific Research Foundation for the introduction of talent (yj2016-4),The Open Fund of State Key Laboratory of Tea Plant Biology and Utilization (SKLTOF20150103).DZ, CW and YW designed the software and website. YY, YZ, YW and YZ initiated these studies. DZ, XC, PW, and CW collected the materials, YY, DZ, YW , GF and YG designed the structure and filled in all data. YY and DZ designed and analyzed data and, with contributions from all authors, wrote the paper. All authors read and approved the final manuscript. We are grateful for stress keywords obtained from the National Joint Engineering Laboratory of crop resistance breeding and disaster reduction.

